# ’Disease-smart’ assisted migration can enhance population fitness and increase resistance to pathogens via immune priming

**DOI:** 10.1101/2024.07.14.603437

**Authors:** Enakshi Ghosh, Matthew Wallace, Ruth A. Hufbauer

## Abstract

1. We studied the potential of combining insect immune priming to synergize with introduction of diverse migrant to safeguard small populations from disease outbreaks that might otherwise lead to extinction.
2. Immune priming in insects refers to the stronger immune response insects have against pathogens after prior exposure. This enhanced immunity can be passed on to offspring and holds promise for insect conservation efforts against diseases.
3. We compared the fitness benefits to a small, inbred population of adding migrants that had not been primed to adding immune primed migrants. While both types of migrants enhanced reproduction, as in cases of genetic rescue, only primed migrants improved survival on exposure to a pathogen.
4. Better immunity led to a trade-off with reproduction in the migrants, but not upon outcrossing with the target population, revealing synergies between hybrid vigor and immune priming.
5. Given the demographic constraints and stochasticity that can exacerbate the effects of disease outbreaks in small populations, combining immune priming with assisted migration offers a proactive strategy to mitigate disease impacts.

## INTRODUCTION

Insects provide vital ecosystem services including pollination, pest control, nutrient transfer, and sustenance for other wildlife, contributing an estimated $577 billion to the world’s economy, annually (Potts et al. 2016). However, the abundance of insects is declining in many areas of the world (Wagner et al. 2021; Harvey et al. 2023). Two key drivers of these declines are habitat loss (along with fragmentation) and disease (Sánchez-Bayo and Wyckhuys 2019).

Habitat loss and fragmentation results in small, isolated populations that are more vulnerable to extinction (Didham et al. 2012). In small populations, genetic drift reduces genetic diversity, which in turn can reduce fitness, increase inbreeding depression, and limit responses to selection imposed by environmental shifts (Willi et al. 2006; Aitken and Whitlock 2013). For instance, isolated populations of butterflies experience inbreeding depression (Nieminen et al. 2001) and even recent habitat loss and fragmentation have been shown to reduce diversity and increase differentiation in insects (Williams et al. 2003). Small populations also suffer from greater demographic stochasticity (Wootton and Pfister 2013), which may amplify these genetic effects (Bensen et al. 2016). In many cases the exact mechanism by which habitat loss and fragmentation suppress populations is unclear, but the outcomes can nonetheless be substantial. For example, habitat loss contributed to a 69% decline in the populations of 45 extant butterfly species (Maes and Van Dyck 2001), coinciding with an increase in the extinction rate from 0.2 to 1.7 species per five years since 1950 (Maes et al. 2022).

Disease caused by parasites and pathogenic microbes have been linked to the decline of many insect species and can cause rapid decreases in population size (Goblirsch 2018; Raymann et al. 2018). For example, high infection rates with *Variomorpha* parasite have been implicated in the rapid decline of several North American bumble bee species, such as *Bombus affinis* and *B. occidentalis*, and the likely extinction of one of them (*B. franklini*) (Williams and Osborne 2009; Cameron et al. 2016). Additionally, the effects of habitat loss, particularly the genetic effects, and of pathogens may interact (Spielman et al. 2004; Gibson and Nguyen 2021). For example, prevalence of a parasitic mite increases in populations of *Bombus muscorum* with lower genetic diversity (Whitehorn et al. 2011). Consequently, diseases can cause a significant population fluctuation which can be particularly exacerbated in small, isolated populations.

Currently, insect conservation efforts primarily rely on habitat management or restoration, which are limited by available habitat and often overlook disease considerations (Duffus et al. 2023). A complementary approach is to introduce migrants from other locations with a goal of reducing isolation of populations and attempt to increase genetic diversity and improve fitness (Ralls et al. 2018). The movement of migrants is typically called assisted migration, and an increase in fitness of the recipient populations when inbreeding depression is alleviated is termed genetic rescue (Hufbauer et al. 2015; Kronenberger et al. 2017; Robinson et al. 2017; Frankham et al. 2019). The practice of genetic rescue has gained increasing attention and validation over the last decade, with many documented cases from plants, fish, birds, reptiles, and mammals (Hogg et al. 2006; Robinson et al. 2017; Frankham et al. 2019). All these studies have emphasized that genetic rescue programs are particularly advantageous for small, declining populations for improving long-term population viability (Frankham 2015, 2016). However, these practices need to incorporate considerations of pathogen outbreaks, as the introduction of migrants could potentially introduce new diseases or exacerbate existing ones, threatening the very populations they aim to save. The unpredictable nature of disease dynamics was underscored by a respiratory tract disease outbreak among translocated desert tortoises in the Mojave Desert, despite rigorous pre- translocation screenings (Sandmeier et al. 2009). Similar disease-driven population declines have been observed in various species including black-footed ferrets, Polynesian tree snails, Hawaiian honeycreepers, monarch butterflies, and bumblebees (Cunningham and Daszak 1998; Pelton et al. 2019; Wilson et al. 2019; Fritts 2020). Moreover, pathogen outbreaks can occur independently of genetic rescue efforts, emphasizing the need for integrating solutions for both pathogen outbreaks and long-term fitness enhancement. Currently, attempts to use gene flow to support declining invertebrates are rare (but see an example with Eltham copper butterfly (Roitman et al. 2017)), so there is a need to develop tailored best practices for insects to ensure resilience and sustainability.

While we know that genetic rescue can help small populations, those populations also must overcome demographic constraints imposed by disease. Here, we tested a genetic rescue plan to enhance the overall fitness of a declining population, including improvements in reproduction and defense against pathogen outbreaks. In a successful rescue program, migrants should originate from populations that are genetically and demographically compatible with the target population, sharing similar habitats and exhibiting minimal genetic differences (Ballou and Foose 1996; Bijlsma et al. 2010). Incorporating just 10-20% of migrants for interbreeding seems adequate to enhance population growth (Ovenden 2013; Durkee et al. 2024). While greater genetic diversity can aid adaptability to diseases, the small population size may not endure the time required for host-pathogen co-evolution (Papkou et al. 2021). It may be impractical to expect a migrant population entirely resistant to diseases. However, insects’ innate immunity provides rapid, yet specific defense mechanisms, allowing quick adaptation to new environments. Their immune responses can be enhanced through genetic and environmental manipulations, increasing survival during relocation and preparing them for future outbreaks. This resilience and adaptability present unique opportunities to enhance assisted migration efforts that may be difficult to achieve in vertebrates. Hence, we propose creating a “disease-smart” genetic rescue program by “immune priming” the migrant population before their integration into the target population for interbreeding. Immune priming, akin to a memory-like function in insects’ immune systems, enables a stronger and faster immune response upon subsequent infection (Cooper and Eleftherianos 2017; Sułek et al. 2021). This process, also observed in vertebrates, involves introducing a sublethal dose of the parasite, an incapacitated agent (e.g., heat killed), or specific molecules from the original pathogen (e.g., lipopolysaccharides). Most importantly, the benefits of immune priming can be transferred to the offspring which makes it promising for insect conservation against diseases (Trauer and Hilker 2013; Schulz et al. 2019).

In this study, we explored the potential of immune priming to work synergistically with genetic rescue as a strategy to enhance fitness of small populations. We used red flour beetles (*Tribolium castaneum*) as our insect model and *Bacillus thuringiensis* as the pathogen threat. We first conducted a proof-of-concept experiment to in which we ask: (1) Can immune priming of parents provide offspring protection from disease? (2) Is there evidence of the potential mechanisms of priming via production of hemocytes or melanization? (3) Are there associated trade-offs to improved immunity in terms of reproduction or development time in the absence of disease? We then conducted an experiment with inbred populations as model targets of genetic rescue. We introduced either no migrants, diverse migrants, or diverse, immune primed migrants to ask: (4) In the context of genetic rescue, can immune priming of migrants protect the following generation from disease? (5) Are there associated trade-offs to disease protection in terms of development time or reproduction? The target inbred populations was derived from an admixed stock population and intentionally subjected to a bottleneck event. This mirrors scenarios where large, genetically diverse populations face sudden anthropogenic pressures, leading to inbreeding depression and reduced growth rates. Given the demographic constraints and stochasticity that can exacerbate the effects of disease outbreaks in small populations, immune priming could be a vital strategy. Additionally, the genetic constraints of small, inbred populations limit their ability to adapt to new pathogens, further highlighting the importance of this approach. In scenarios where the resistant migrants aren’t accessible, immune priming could serve as an alternative method to create a resilient migrant population. This could safeguard a small, inbred target population and lay the foundation for a ’disease-smart’ genetic rescue initiative.

## MATERIALS AND METHODS

### Study system and creation of the target population and migrants

Our populations were derived from a genetically diverse stock comprising five lineages of *T. castaneum* (Durkee et al. 2024). The stock population was maintained at 2000 individuals housed in replicate enclosures 4cm × 4cm × 6cm containing 100 mL of medium (95% wheat flour, 5% brewer’s yeast), at 31°C and 54 ± 10% relative humidity. The target population was one of the populations from Durkee et al. (Durkee et al. 2024). Briefly, the target population was founded with 50 individuals, thus imposing a mild demographic bottleneck. It then was reared in a challenging environment that had deltamethrin, an insecticide added to the medium for eight generations. This event simulated a sudden environmental change, and imposed a selective bottleneck. At the end of the Durkee et al. (2024) experiment, the population was maintained on standard media (without pesticide) for seven additional generations during which population size was at least 500 individuals. This history of a demographic bottleneck and strong selection, with the environment changing to more than less challenging conditions is not unlike what populations experience in nature, and provides a target population that should have reduced genetic variation and so could benefit from assisted migration.

Our migrants were derived from the genetically diverse stock population described above. In preparation for our experiment, we selected 50 individuals from this population and placed them in 4 × 4 × 6-cm enclosures partially filled with 30 mL of flour and yeast medium for thirteen generations. This subset represented an isolated healthy population which showed no inbreeding depression (monitored though growth rate).

To create primed migrants, we separated female pupae from the migrant pool to develop to adulthood for priming as unmated females. We focused on females (“maternal priming”) for several reasons. First, females are often preferred in genetic rescue programs because they can benefit small populations by introducing mitochondrial DNA (Tallmon et al. 2004) Second, previous evidence shows that maternal priming can have trans-generational effects in insects (Little et al. 2003; Sadd and Schmid-Hempel 2007; Freitak et al. 2009; Zanchi et al. 2011). Third, red flour beetles’ are polyandrous, and thus males have highly variable reproductive success while females are more consistent.

*Bacillus thuringiensis* culture was prepared from a −80 °C glycerol stock, cultured overnight in nutrient broth at 30°C until reaching an optical density of 0.95 at 600nm. After centrifugation, the pellet was resuspended in 1mL PBS. Bacteria were either heat-killed for priming or used live for larval infections. Heat-killed bacteria were adjusted to 10^11^cells/mL of PBS. We ensured this step by plating the heat-killed bacteria on agar plates and incubated them overnight for confirmation. For priming, adult females were pricked between the head and thorax with a 0.1-mm pin dipped in the heat-killed bacterial solution. A sham control was created using PBS for pricking. Pricked females were allowed to heal for 48h before mating.

### Experiment 1: Priming confirmation and evaluation of potential trade-offs

To confirm that maternal (G_0_) immune priming can provide survival benefits to the next generation (G_1_) against the pathogen, we created three sub-populations for comparison. 1) Unprimed: 50 beetles (with a 1:1 male-to-female ratio) housed in a 4 × 4 × 6-cm enclosure, 2) Sham primed: 25 sham-pricked females housed with 25 control males in a patch, and 3) Primed: 25 primed females (pricked with heat-killed bacterial solution) housed with 25 control males in a patch (n=30 patches, 50 beetles/patch).

After a 48-hour oviposition period, we sieved the adults from the media, retaining the eggs (G_1_), which were then maintained in the same media until they reached the larval stage. These larvae (17 days post oviposition) were divided into four subsets to evaluate the effects of maternal priming on 1) larval survival at 24-, 48-, and 72-hours following exposure to the Bt pathogen and the percentage of challenged larvae successfully developed into adults, 2) immunological function, 3) offspring production per mating pair, and 4) development duration and adult body weight.

To evaluate survival and formation of adults in the presence of Bt, 150 larvae per treatment were exposed to a lethal dose of the bacterium, and survival was evaluated at 24, 48, and 72 hours post-exposure, and the percentage of larvae successfully reaching the adult stage.

Immunological function was assessed on 50 larvae per treatment by measuring the concentration of hemocytes and the production of melanin following previously established protocols (Ghosh et al. 2023a, b). Briefly, larvae were incised laterally, and hemolymph was collected into pre-chilled Eppendorf tubes. To count hemocytes, hemolymph was diluted 1:2 with PBS, and 5 μL of this mixture was loaded onto a Neubauer hemocytometer, then counted under a microscope (200x magnification) after 15 minutes. We used phenoloxidase activity to assesse production of melanin by centrifuging hemolymph (1:2 with ice-cold PBS), adding the supernatant to a microplate well with 2 mM L-Dopa (substrate), and recording melanization spectrophotometrically at 490 nm after 30 minutes (n = 20/treatment).

To evaluate if maternal immune priming leads to potential trade-offs with reproduction in G_1_ progeny, we set up single mating pairs (21-30 pairs/treatment) in plastic vials containing 5g of flour. After a 48-hour oviposition period, we sieved the adults. Offspring from each pair were counted on day 35. To evaluate potential trade-offs with other life-history traits the duration of egg-to-pupae development in days and the body weight of adult beetles were recorded on 50 larvae per treatment.

### Experiment 2: Integrating primed migrants into genetic rescue

In our genetic rescue experiment, we added diverse migrants to the target population, which given some expected reduction in diversity in that population should increase fitness (by masking deleterious mutations that lead to genetic load), as is typical in efforts at genetic rescue. Our goal was then to examine whether immune priming could provide further benefits to the target population, through enhancing survival in the presence of a pathogen. As such, we implemented three treatments: (1) the target population without any migrants, serving as the control, (2) the target population with migrants that had not been immune primed (unprimed migrants), (3) the target population with primed migrants.

Replicate control populations were initiated with 50 beetles from the target population stock. Populations receiving migrants (of either type) were initiated with 40 individuals from the target stock, to which 10 unmated migrant females were added (20% of the population). This approach ensures that the effects of migrants are primarily genetic rather than demographic (Hufbauer et al. 2015). Although the ideal percentage of migrants used by managers may depend upon population size, existing genetic diversity, and the specific objectives of the initiative, typically, genetic rescue endeavors, including translocation programs, incorporate an introduction of migrants ranging from 10% to 20% of the recipient population (Hogg et al. 2006; Whiteley et al. 2015).

These experimental groups were given 48 hours for mating and oviposition, after which adults were removed. Larvae were collected at 17 days post-oviposition and subdivided into three groups as above to assess 1) larval survival at 24-, 48-, and 72-hours following exposure to the Bt pathogen, 2) percentage of challenged larvae successfully developed into adults, 3) offspring production per mating pair, and 4) development duration.

### Statistical analysis

Our first experiment aimed to assess whether trans-generational immune priming can improve survival of offspring exposed to a pathogen, as well as associated trade-offs. To accomplish this, we used a log-rank model to compare larval survival among control, sham, and primed migrants, after ensuring the proportional hazards assumption held true. We analyzed percent adult formation and adult body weight data using a generalized linear model, followed by Tukey’s multiple comparison test. Since residuals from offspring production and development duration deviated from the normal distribution (confirmed by Shapiro-Wilks test), we used the Kruskal-Wallis test with Dunn’s multiple comparison. Immunological tests, total hemocyte count and degree of melanization, were assessed using the non-parametric Mann-Whitney U test.

Our second experiment aimed to test if primed migrants can improve survival of individuals from the target population during a pathogen outbreak, while still offering benefits from outcrossing. Accordingly, we compared larval survival between the control populations, and the populations that received unprimed migrants and primed migrants using a similar log-rank model as described above. Data on adult formation, development duration, offspring production were analyzed using the Kruskal-Wallis test with Dunn’s multiple comparison. All statistical analyses were performed using GraphPad Prism software.

## RESULTS

### Immune priming of parents protects offspring from disease

The efficacy of trans-generational immune priming was evaluated through two key metrics: the probability of survival upon exposure to a lethal pathogen (Fig. 1a) and subsequent formation of adults (Fig. 1b). We found that maternal priming enhanced survival among the G_1_ progeny (Log-rank model: X^2^= 54.44, *p* < 0.0001). Upon challenging them with the pathogen, a stark contrast was observed in mortality rates within 48 hours. Unprimed and Sham primed individuals experienced mortality rates of 76-77%, whereas only 24% of Primed migrants succumbed to the challenge. Furthermore, the ability of challenged individuals to transition into adulthood varied significantly. Completion of development to adulthood tracks survival. Only 12-15% of Unprimed and Sham primed individuals reached adulthood while 60% of Primed individuals eclosed successfully as adults (GLM: F_2_= 339, *p* < 0.0001). No notable differences were observed between the Unprimed and Sham primed groups (GLM: F_2_= 339, *p* = 0.62), suggesting that the priming method involving pricking adult beetles did not impose any survival challenges.

**Figure 1:**
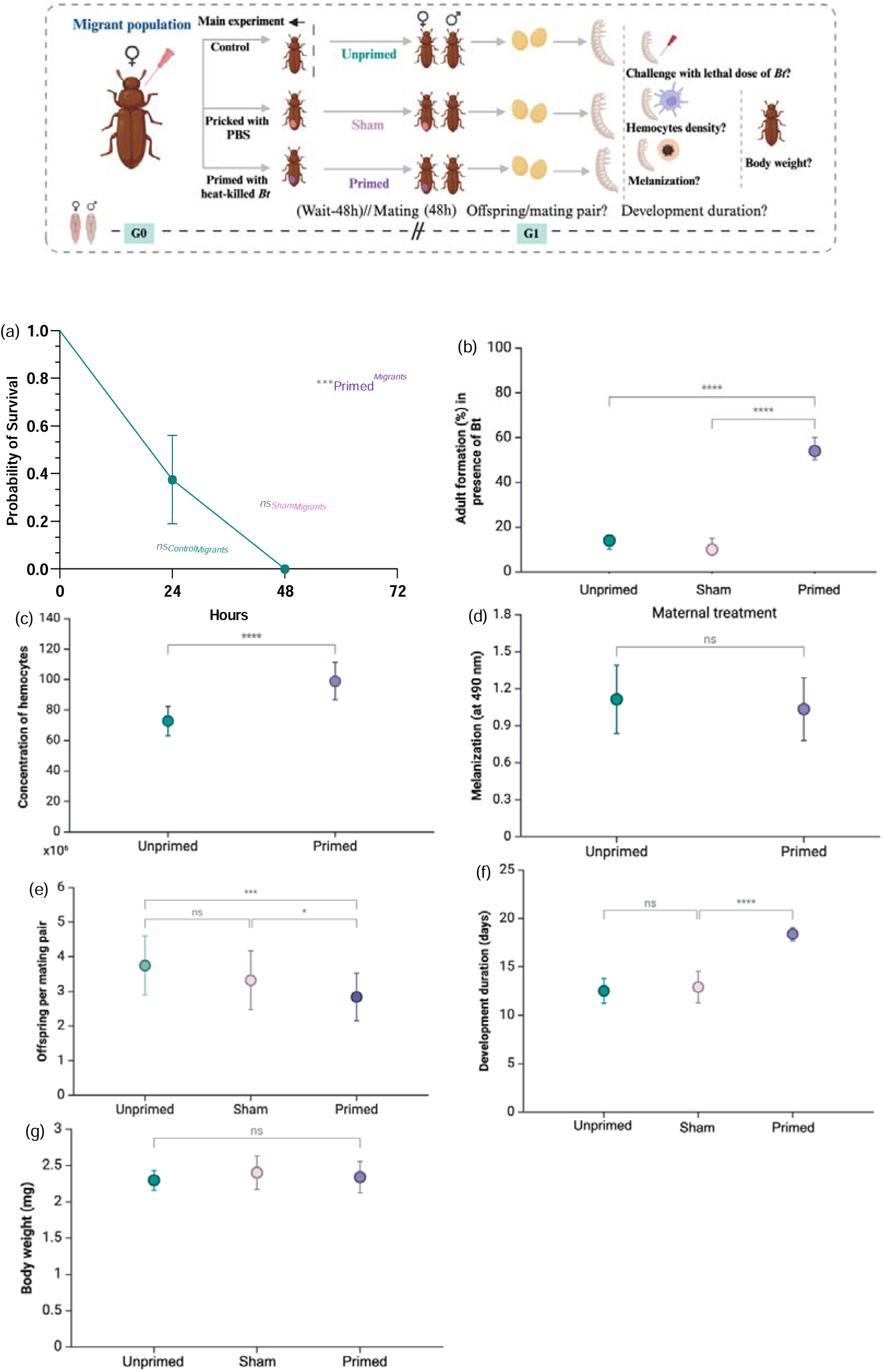
Impact of maternal immune priming on progeny. In the presence of a pathogen *(Bt*) (a) progeny survival increases with priming and (b) the percent of progeny reaching adulthood increases with priming. Mechanistically, the increased immunity coincides with (c) increased density of hemocytes in larvae (d) but not with evident changes in melanization. Performance in the absence of the pathogen shows (e) reduced offspring per mating pair and (f) increased development time in immune primed individuals, but (g) no clear shift in body weight. Data represents mean with 95% confidence intervals, statistical significance is based on *p-*values.

### Enhanced survival coincides with increased production of hemocytes

The concentration of hemocytes in Primed larvae registered 1.4 times higher than the Unprimed group (Fig. 1c, *U*= 68.5, *p* < 0.0001). This indicates significant enhancement in basal cellular immunity conferred by trans-generational immune priming. Conversely, our assessment of melanization capacity revealed considerable variability within each group of individuals, yielding no significant difference in the treatment groups (Fig. 1d, *U*= 178.5, *p*= 0.498).

### Enhanced survival and improved immunity show trade-offs with reproduction and larval development duration

To evaluate potential costs of priming, we assessed offspring production per mating pair in the absence of disease. Primed females laid fewer eggs/day than Unprimed and Sham primed females (Fig. 1e). Unprimed females averaged four eggs per day, whereas Primed females produced two to three eggs daily (*U*= 14.18, *p* < 0.0001). Furthermore, maternal immune priming increased the time it took for larvae to complete development from 12-13 days for Unprimed and Sham primed individuals to 18 days in Primed individuals (Fig. 1f, *U*= 88.64, *p* < 0.0001). In contrast to reproduction and development time, weight of adult beetles revealed no significant difference between Unprimed, sham primed and Primed migrant groups (Fig. 1g; GLM: F_2_ = 2.02, *p* = 0.136).

### Introducing primed migrants in the context of genetic rescue provides dual fitness benefits to the target population

Our second experiment aimed to evaluate whether introducing primed migrants could enhance the survival of the target population during a pathogen outbreak. Upon subjecting the populations to a pathogen challenge, the target populations receiving primed migrants exhibited a threefold higher survival compared to Unprimed migrants. In contrast, we observed no clear difference in the probability of survival between the target populations receiving no migrants and those receiving Unprimed migrants, indicating that increased genetic diversity alone does not account for this survival advantage (Fig. 2a, X^2^= 114.9, *p* < 0.0001). Furthermore, ∼40% more adults emerged from populations receiving Primed than Control migrants (Fig. 2b, *U*= 10.52, *p* < 0.0001).

**Figure 2:**
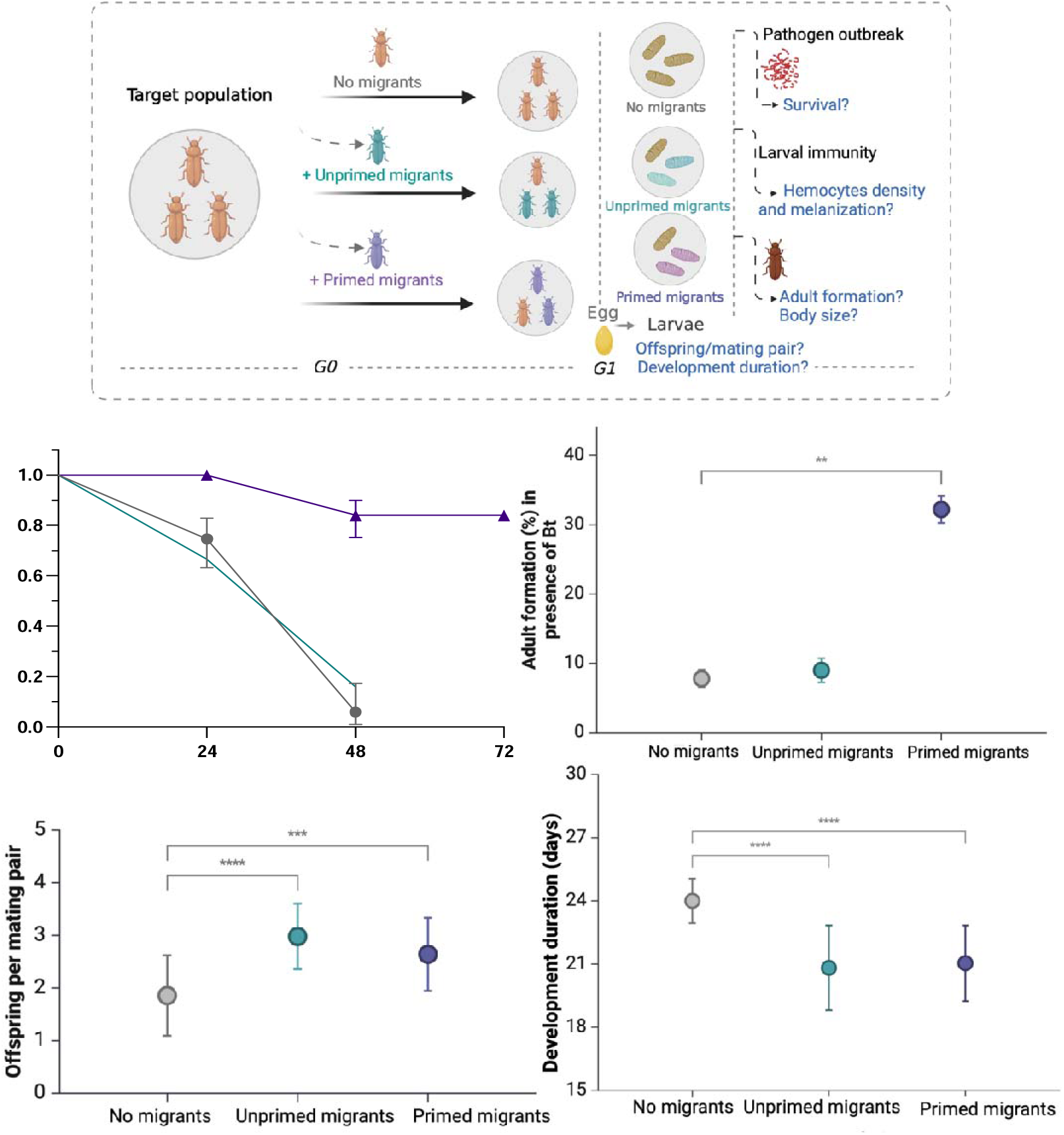
Impact of introducing primed migrants into the target population: In the presence of a pathogen *(Bt*), (a) progeny survival increases and (b) the percent of progeny reaching adulthood increases. Performance in the absence of the pathogen shows no trade-offs associated with (c) offspring production per mating pair and (f) development duration. Data represents mean with 95% confidence intervals, statistical significance is based on *p-*values.

We further asked if introducing primed migrants were comparable to unprimed migrants in being able to increase reproductive success in an inbred target population, while also providing additional benefits in during a pathogen outbreak. To achieve this, we evaluated reproduction of the target population by quantifying the number of offspring produced the generation following migrant addition. Our analysis revealed that both types of migrants, Unprimed and Primed, increased offspring production (Fig. 2c, *U*= 14.18, *p*< 0.0001). Importantly, there were no significant differences between Unprimed and Primed migrants (*U*= 14.18, *p* = 0.1239). This indicates that, even though the enhanced immunity due to priming led to reduced reproduction without outcrossing (Figure 1e), the introduction of primed migrants to did not, and thus the benefit of outcrossing with the diverse migrants was more potent than the potential cost of individual-level enhanced immunity. Additionally, the introduction of both Primed and Unprimed migrants reduced development time of their target populations by 3-4 days (Fig. 2d, *U*= 63.76, *p* < 0.0001).

## DISCUSSION

Our findings indicate that the introduction of immune-primed migrants not only enhances reproduction, effecting genetic rescue, but also fortifies the population against future disease outbreaks. We compared two types of migrants to enhance the survival of a target population in the presence and absence of a pathogen. The introduction of migrants aimed at adding genetic diversity resulted in notable improvements in fitness, primarily evidenced by increased offspring production, thereby reducing the extinction risk of the target population. However, augmenting genetic diversity was not sufficient to shield the population from pathogen outbreaks. While genetic diversity can bolster declining populations and aid in their conservation efforts, the sudden onset of pathogen outbreaks poses a significant threat to the success of such rescue plans. To address this concern, we introduced primed migrants selected for their defense benefits. This served two primary purposes. Firstly, primed migrants were introduced in anticipation of pathogen outbreaks, effectively preempting potential threats. Secondly, these migrants served as proxies for a naturally resistant population, offering an alternative solution when establishing a resistant population directly may not be feasible.

### Integration of insect immune priming in conservation efforts

Insects possess an innate immune system devoid of the pathogen-specific antibodies and memory cells that are hallmarks of vertebrate acquired immunity, yet they exhibit remarkable efficiency in eradicating foreign invaders (Sułek et al. 2021). This efficacy is partly due to immune priming, which is akin to acquired immunity observed in vertebrates (Cooper and Eleftherianos 2017). This process functions as a form of memory where recognition of re- introduced microbes (or part of it) induces a faster and higher immune reaction leading to elimination or reduction of the invaders. While distinct from conventional vaccines, immune priming confers analogous benefits and can be conceptualized as an insect-specific form of vaccination. Notably, this approach has been successfully implemented to protect *Apis mellifera* colonies against American foulbrood disease (2023). In our investigation, we asked whether immune priming could improve insect conservation within the context of genetic rescue program.

Delivering heat-killed bacteria to our migrant populations bolstered disease resistance in their offspring by enhancing cellular immunity. These data match previous work of immune priming across diverse insect taxa, including beetles, moths, butterflies, bees, flies, mosquitoes, and bugs (Sadd and Schmid-Hempel 2007; Freitak et al. 2009; Sheehan et al. 2020; Sułek et al. 2021; Li et al. 2022; Gálvez et al. 2024). Although the precise mechanisms underlying immune priming and their transgenerational effect are not fully understood, it might involve epigenetic alterations, particularly histone modifications, which facilitate heightened gene transcription upon subsequent exposure to pathogens (Sułek et al. 2021). Recent studies in honeybees (*Apis mellifera*) have demonstrated that the transmission of immune signals from mother to offspring occurs through the direct transfer of bacterial components that bind to the egg-yolk protein vitellogenin (Vilcinskas 2021). This transmission enhances resistance to pathogens in subsequent generations. In our experiment, priming mediated defense was subsequently transmitted to the target population as they bred with primed migrant females. Introducing only ten primed migrant individuals resulted in a threefold increase in survival compared to unprimed migrant treatments. Additionally, the observed rise in offspring production following the rescue effort highlights its effectiveness in augmenting genetic variation and mitigating inbreeding depression. Increased genetic diversity also enhances the adaptive potential of the population to environmental changes (Ballou and Foose 1996). However, the time needed for a small population to adapt to pathogen pressure increases the risk of extinction. This risk can be minimized by considering disease while implementing conservation efforts like genetic rescue where migrants can be vaccinated or primed before introducing to the target population. The decision to prime the migrants instead of the target population under focus can be justified by several key reasons. Firstly, there are observed trade-offs between heightened protection and reproduction, making it essential to prioritize immune priming where it can be most effective. Secondly, migrants may face increased susceptibility to pathogen outbreaks due to factors such as stress during migration or exposure to novel pathogens in new environments. Lastly, logistical considerations may favor the implementation of immune priming in migrants, where accessibility and control may be more feasible.

*Benefits vs limitation* (change this sub-title? Synergies Between Hybrid Vigor and Immune Priming?)

Immune priming often involves a trade-off with reproductive success. For instance, in species like *Manduca sexta* and mosquitoes, offspring of primed parents tend to lay fewer eggs compared to those from control parents, indicating a trade-off between immune priming and egg production (Adamo et al. 2023; Cime-Castillo et al. 2023). We observed this trade-off in our study using *Tribolium* as well, primed females produced fewer offspring than their unprimed counterparts. This pattern is consistent with the classic resource allocation model, which suggest that competition for limited resources precludes high reproduction when immunity is upregulated. This trade-off can manifest at the physiological level (within an individual) as seen here, and at the evolutionary level (genetic distinction among individuals in a population) (Schwenke et al. 2016). Although these findings present a present a potential drawback to using immune priming in conservation, our investigation revealed a promising shift: when primed migrants mated with the target population, this trade-off disappeared.

Notably, reproduction of populations receiving unprimed and primed migrants did not differ significantly, and both groups produced more offspring than the target population.

We chose a target population likely to have low genetic diversity, as these are the targets of assisted gene flow in nature. It had passed through a selective bottleneck and been maintained in the laboratory for multiple generations following that bottleneck. The introduction of migrants from a genetically diverse population provided the opportunity for outcrossing, and the production of offspring with increased fitness compared to the original population, thus elevating demographic vital rates. This could result from either heterosis, where hybrid vigor enhances traits, or adaptive evolution, which favors newly introduced or recombinant genotypes better suited to the environment, rapidly improving population vital rates (Whiteley et al. 2015). The disappearance of the trade-off between reproduction and immunity in hybrids may be attributed to intrinsic selection, allowing the expression of beneficial genetic traits, or extrinsic selection, which pertains to environment-specific fitness, or a combination of both factors (Burke and Arnold 2001). Nonetheless, it’s important to note that we assessed reproductive success by counting egg laying within a 48-hour period only. It’s possible that there’s a trade-off with their lifetime egg laying or with other life-history traits which warrants further investigation.

### Towards creating a disease smart insect conservation plan

Insect populations face unpredictable impacts from anthropogenic factors, including disease outbreaks (Wagner et al. 2021). To mitigate these risks, conservation efforts must adopt an anticipatory approach by integrating disease considerations early in the process. We provide an example of how within a genetic rescue program, the concept of immune priming can be integrated to have dual benefits of higher reproduction and protection against disease. This approach represents a significant advance beyond current practices, which often rely solely on health evaluations and quarantine measures to mitigate disease outbreaks, yet these measures may not be entirely effective in preventing all disease occurrences. Unforeseen impacts on behavior and physiology in new environments can heighten disease susceptibility or transmission risks, underscoring the need for practitioners to anticipate and mitigate disease impacts during translocation efforts.

## Authors’ contributions

E.G. conceived the idea, developed the methods, designed the study, conducted the study, analysed the data, made the figures and wrote the manuscript. M.W. assisted with conducting the study and with writing. R.A.H. assisted with conceiving the idea and study design, as well as writing the manuscript, and providing funding.

## Acknowledgements

This work was supported by the U.S. National Science Foundation DEB-1930650 and by the USDA NIFA (Hatch Project 1012868) to RAH.

